# Cheese or cheese infusions – ecological traps for mosquitoes and spotted wing Drosophila

**DOI:** 10.1101/2020.12.30.424879

**Authors:** Daniel Peach, Max Almond, Elton Ko, Sanam Meraj, Regine Gries, Gerhard Gries

## Abstract

We tested the hypothesis that the “ecological trap” phenomenon (a mismatch between a habitat’s perceived attractiveness and its actual quality, resulting in a population sink) is exploitable for pest control. We selected mosquitoes as modal organisms, because selection of an oviposition site by adult female mosquitoes in response to its perceived attractiveness is of paramount importance for the development and survival of their larval offspring. In laboratory and/or field experiments, we show that (*i*) each of five cheese varieties tested (Raclette, Pecorino, Brie, Gruyere, Limburger) strongly attracts females of both the yellow fever mosquito, *Aedes aegypti*, and the common house mosquito, *Cx. pipiens;* (*ii*) cheese infusions, or headspace odorant extracts (HOEs) of cheese infusions, significantly affect oviposition choices by *Cx. pipiens* and *Ae. aegypti*, (*iii*) HOEs contain at least 13 odorants; (*iv*) in field settings, cheese infusions more effectively stimulate oviposition by *Cx. pipiens* and *Culiseta incidens* than bluegrass (*Poa* sp.) infusions, and also capture (by drowning) the spotted wing *Drosophila* (SWD); (*v*) the microbe composition of home-made cheese infusions modulates oviposition choices by mosquitoes; and (*vi*) the type of cheese infusion coupled with its nutritional content strongly affects the survivorship of mosquito larvae. In combination, our data show that microbial metabolites associated with cheese and cheese infusions are both attractive to adult mosquitoes seeking hosts and oviposition sites, respectively, and are toxic to mosquito larvae. These microbes and their metabolites could thus be coopted for both the attract and the kill function of “attract & kill” mosquito control tactics. Implementation of customizable and non-conventional nutritional media, such as home-made cheese infusions, as microbe-based ecological traps presents a promising concept which exploits insect ecology for insect control.

## Introduction

Insects have profound impact on humans. Locust swarms of biblical proportions threaten famine in several countries [1, 2], declining pollinator populations put the production of agricultural produce in jeopardy [3, 4], range expansions of pest insects incur massive global costs [5, 6], and insect vectors of pathogens account for debilitating diseases or even death of millions of humans every year [7]. In response, humans have dedicated immense amounts of resources and time to manipulating pest insects. However, with increasing development of pesticide resistance [8,9], environmental concerns about the use of pesticides [10, 11], public pushback against the deployment of genetic tools [12], and the scarcity of new insect control tools being developed and becoming widely available [13], innovative approaches to the management and control of pest insects are desperately needed.

Harnessing insect ecology for insect control is an innovative concept proposed for many of the World’s most impactful pest organisms, including the medically important vectors of deadly pathogens [14–16] and the many plant herbivores that threaten food security [17, 18]. The concept seeks to exploit, among others, the ecological interactions between insects and microbes for improved control of pest insects.

Microbes are cosmopolitan in the environment, coming into contact and interacting with animals in their habitat [19], on their body [20] and on their food [21, 22], often residing in the internal environment of an animal itself, both intra- and extra-cellularly [23, 24]. Animal-microbe associations are ancient, and insects, being supremely speciose [25], have formed myriad ecological interactions with microbes [26–28]. Insects disperse microbes between habitats [29], serve as microbe hosts [22, 29], and – in some cases – even farm microbes [30]. Microbes, in turn, affect insect nutrition, immune response, and many types of insect behaviour [20, 22, 26–28, 31]. For example, microbe-derived semiochemicals guide foraging insects to food or oviposition resources which can also serve as suitable habitat for microbes [22, 26, 32]. Microbe-derived semiochemicals contribute to the attraction of flies to food [33], yellowjackets to fruit [34], pollinators to flowers [31], mosquitoes to vertebrate hosts [35] and oviposition sites [36], and the attraction of aphid predators and parasitoids, as well as mosquitoes, to aphid honeydew [37, 38].

Microbe-derived semiochemicals may not always accurately convey the quality of a resource to foraging insects, possibly resulting in an ecological trap, a mismatch between a habitat’s perceived attractiveness and its actual quality, causing a population sink [39–41]. Mismatches may occur when, e.g., insects respond to microbe odorants from a habitat generalist, rather than a specialist, currently residing in a habitat unsuitable for the foraging insect or its future offspring. Mosquitoes (Diptera: Culicidae) use microbe-derived semiochemicals as resource indicators in many parts of their complex life history including host-foraging and ovipostion site selection [35, 36]. Female mosquitoes do not exhibit parental care and select biotypes as oviposition sites that meet a select range of biotic and abiotic conditions suitable for the development and survival of their larval offspring. Female mosquitoes assess biotype characteristics such as salinity [42], predator presence [43], and microbial abundance and composition [36] based, in part, on microbe-derived semiochemicals [36, 44, 45] and discern oviposition sites accordingly. Similarly, mosquitoes exploit the odorants from human skin microbiota to locate human hosts [20, 35, 46, 47] or substances such as Limburger cheese on which similar microbes can grow [48–50]. However, microbe odorants can also misguide resource-foraging mosquitoes. For example, decomposing foliar material of common blackberry, *Rubus allegheniensis*, in oviposition sites of *Culex pipiens*, attracts adult females for oviposition but kill larval offspring [41]. Similarly, blood-seeking female *Anopheles* and *Aedes* mosquitoes responding to the microbial odor of Limburger cheese reach a resource from which they cannot feed [49–51].

Odorants from cheese and its microbes have been implicated in various insects’ attraction to cheese. For instance, butyric acid [52] and possibly microbial odorants [53] attract the aptly-named cheese skipper, *Piophila casei*, to cheese as an oviposition resource [54]. Similarly, *An. gambiae* mosquitoes respond to the odor of Limburger cheese which resembles that of human feet [55]. The odor is likely produced by *Brevibacterium linens* [56], a bacterium (i) involved in the ripening of Limburger cheese, (ii) related to microbes that are thought to cause the “smelly foot” odor of Limburger cheese, and (iii) closely related to microbes found on human feet [49, 56].

Microbes are essential ingredients for cheesemaking and are responsible for the many distinctive flavours, odours, and textures that are characteristic of certain types of cheese [57–60]. Some microbes that are co-opted for cheesemaking also occur in the microbiome on human skin [56] and in either context may produce the same odorants that attract mosquitoes. *Lactobacillus* spp. exemplify microbes that occur in many contexts or ecological niches. Members of this genus are used in cheesemaking [61], colonize human skin [62], and reside in inflorescences [63]. Some *Lactobacillus* spp. produce lactic acid [64] which is an important mosquito attractant [65]. *Lactococcus lactis*, another lactic acid-producing bacterium used for cheesemaking [66], is one of several microbes that may attract mosquitoes to oviposition sites [36].

Here we demonstrate that inter-kingdom signaling between microbes and mosquitoes, as well microbes and the spotted wing Drosophila (SWD), *Drosophila suzukii* (a major agricultural pest), can be harnessed for the atttraction and control of mosquitoes and SWD. We show that SWD and two medically important mosquitoes, Aedes aegypti and *Culex pipiens*, are strongly attracted to cheese or home-made cheese infusions which become ecological traps, killing the adult insects or their larval offspring. Furthermore, we investigate the mechanisms underlying the attractiveness of these ecological traps and promote their use as an affordable and innovative means of pest control.

## Results

### Cheese as an attractant to host-seeking mosquitoes in the laboratory

To test the attractiveness of cheese to host-seeking mosquitoes, we deployed paired, adhesive-coated traps inside mesh cages housing Ae. aegypti or *Cx. pipiens* (Figure 1A) and baited one trap in each pair with 20 g of cheese. Each of five cheese varieties tested (Raclette, Pecorino, Brie, Gruyere, Limburger) equally and strongly attracted females of both Ae. aegypti and Cx. pipiens (Figure 1B; Exp. 1: n = 10, z = 10.0, p < 0.0001; Exp. 2: n = 5, z = 4.3, p < 0.0001; Exp. 3: n = 10, z = 7.1, p < 0.0001; Exp. 4: n = 10, z = 6.9, p < 0.0001; Exp. 5: n = 10, z = 11.5, p < 0.0001; Exp. 6: n = 9, z = 11.9, p < 0.0001; Exp. 7: n = 11, z = 10.6, p < 0.0001; Exp. 8: n = 10, z = 12.1, p < 0.0001; Exp. 9: n = 10, z = 10.3, p < 0.0001; Exp. 10: n = 6, z = 6.1, p < 0.0001).

**Fig. 1.**
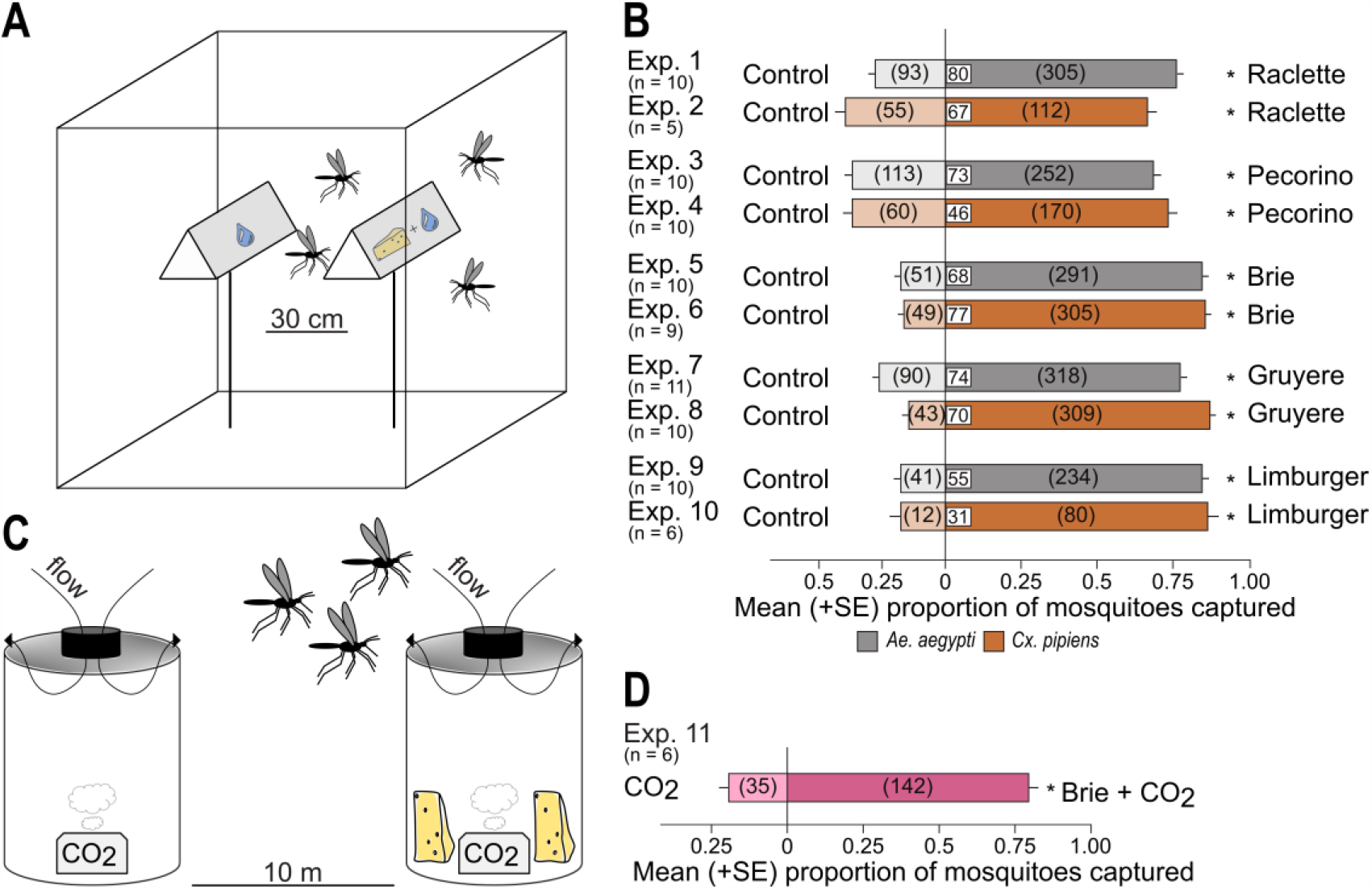
Effect of cheese on attraction of mosquitoes in laboratory and field experiments. **(A)** Graphical illustration of the paired-trap experimental design used in laboratory experiments 1-10; **(B**) Effect of cheese variety on attraction and capture of 1-to 3-day-old female *Aedes aegypti* and *Culex pipiens* in laboratory experiments 1-10; **(C**) Graphical illustration of traps (Biogents AG, Regensburg, Germany) and baits deployed in field settings (Exp. 11); and **(D**) Comparative captures of mosquitoes (primarily *Culiseta incidens* and *Cx. pipiens*) in field traps baited with CO2 alone and in binary combination with Brie cheese. An asterisk indicates significant preference (p < 0.05) for the specific test stimulus (binary logistic regression analysis with logit link function); numbers within white boxes in bars indicate the mean proportion of mosquitoes captured, and numbers within brackets indicate the total number of mosquitoes captured in traps baited the respective stimulus.

#### Cheese as an attractant to host-seeking mosquitoes in field settings

To test the attractiveness of cheese to host-seeking wild mosquitoes in field settings, we used paired BG sentinel traps (Figure 1C) and baited one trap in each pair with CO2 and the other with CO2 and Brie cheese (20 g). The presence of cheese in trap baits afforded significantly greater captures of mosquitoes (n = 6, z = 7.42, p < 0.000; Figure 1D), mainly *Culiseta incidens* and *Cx. pipiens*.

#### Effect of cheese infusions as oviposition stimulants for mosquitoes (laboratory experiments)

To test the effect of cheese infusions as oviposition stimulants for mosquitoes, we prepared cheese infusions by fermenting 50 g of cheese in 5 L of water for 5 days in a rain-sheltered area outdoors, subjecting plain water as a control stimulus to the same protocol. In each experimental replicate, we then offered 20 gravid female *Cx. pipiens* or female *Ae. aegypti* a choice between four cups filled with 200 mL of a cheese infusion or filled with plain water (Figure 2A), recording their selection of oviposition site 48 h later. Cheese infusions had a significant effect on oviposition behaviour of *Cx. pipiens* females (n = 10, χ2 = 33.48, p <0.0001; Figure 2B), with Brie and Raclette cheese infusions receiving the same number of egg rafts (z = −0.76, p = 0.45) but significantly more egg rafts than Limburger cheese infusions (Brie: z = 4.87, p < 0.0001; Raclette: z = 5.42, p < 0.0001) or than plain water (Brie: z = 2.02, p = 0.04; Raclette: z = 2.76, p = 0.006). Tested head-to-head, the Brie infusion received more egg rafts than the Raclette infusion (n = 10, z = 3.95, p < 0.0001; Figure 2C).

**Fig. 2.**
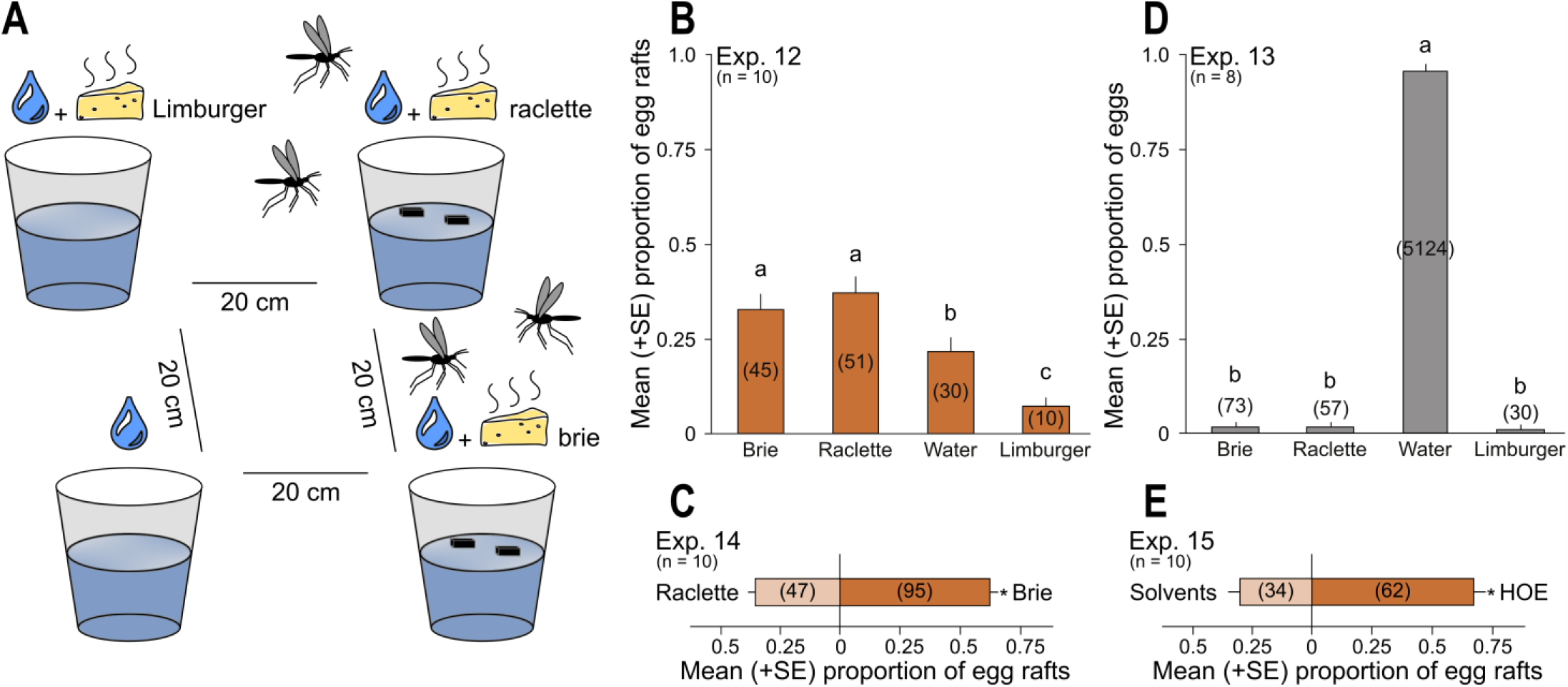
Cheese infusions as oviposition cues for gravid mosquitoes. (**A**) Graphical illustration of the experimental design applied in **B, C** and **D** (with only two, instead of four, cups used in **C**). **(B, D**) Effect of cheese variety in cheese infusions on oviposition preference by groups of 20 female *Culex pipiens* each (n = 10) **(B**) and 20 female *Aedes aegypti* each (n = 8) **(D). (C)** Effect of Brie cheese infusion, or Raclette cheese infusion, on oviposition preference by groups of 20 *Cx. pipiens* each (n = 10). **(E)** Effect of headspace odorant extract (HOE) of Brie cheese infusion on oviposition preference by groups of 10 female *Cx. pipiens* each (n = 10). All mosquitoes were 8-15 days old and 4-5 days post blood-feeding. Different letters on bars within experiments indicate differences in the mean proportion of mosquitoes responding to respective stimuli (type 3 analysis of effects with Tukey’s HSD, p < 0.05). An asterisk indicates a significant preference (p < 0.05) for the specific test stimulus (binary logistic regression analysis with logit link function). Numbers within brackets indicate the total number of eggs or egg rafts oviposited in response to the test stimulus.

Cheese infusions also had a significant effect on oviposition behaviour of Ae. aegypti females (n = 8, χ2 = 123.2, p <0.0001; Figure 2D). Water received more Ae. aegypti eggs than Raclette infusions (z = 8.4, p < 0.0001), Brie infusions (z = 8.3, p < 0.0001), or Limburger infusions (z = 7.4, p < 0.0001), with all cheese infusions being equally deterrent (Brie-Raclette: z = −0.08, p = 0.94; Brie-Limburger: z = 0.42, p = 0.68; Raclette-Limburger: z = 0.5, p = 0.62).

### Identification of odorants associated with cheese infusions

To identify the cheese odorants that may mediate oviposition behavior by mosquitoes, we (i) collected headspace odorants of Brie cheese infusion on Porapak-Q™ adsorbent, (ii) desorbed the odorants with solvent, and (iii) offered female Cx. pipiens a choice between oviposition sites conjoined with aliquots of headspace odorant extract (HOE) or a solvent control. When oviposition sites conjoined with HOE extract received significantly more egg rafts (n = 10, z = 3.3, p = 0.001; Figure 2E), we proceeded with odorant identifications by gas chromatography-mass spectrometry (GC-MS). Analyzing HOEs not only of Brie cheese infusions (Figure 3A, Table 1), but also of Raclette cheese infusions (Figure 3B, Table 1), and of Limburger cheese infusion (Figure 3C, Table 1), we identified a total of 13 odorants (1-hexanol; dimethyl trisulfide; phenol; p-cresol; nonanal; decanal; indole; 3-methylbutyric acid; 2-methylbutyic acid; hexanoic acid; benzaldehyde; S-methyl hexanethioate; S-methyl octanetioate), with significant odorant overlap between sources.

**Table 1.** Headspace odorants collected from a Brie, Raclette, or Limburger cheese infusion (see methods for details.

| Compound | % total |  |  |
| --- | --- | --- | --- |
|  | Brie | Raclette | Limburger |
| 3-methylbutyric acid | – | 1.6 | 2.7 |
| 2-methylbutyric acid | – | 2.2 | 3.7 |
| 1-hexanol | 0.8 | – | – |
| hexanoic acid | – | – | 2.2 |
| dimethyl trisulfide | 3.5 | 1.2 | 2.6 |
| phenol | 23.6 | 51.6 | 50 |
| benzaldehyde | – | – | 2.8 |
| p-cresol | 48.4 | 25.0 | 15.4 |
| S-methyl hexanethioate | – | – | 0.7 |
| nonanal | 2.1 | 1.2 | 0.8 |
| decanal | 1.5 | 0.8 | 0.3 |
| indole | 8.7 | 12.7 | 13.1 |
| S-methyl octanethioate | – | – | 0.7 |

**Fig. 3.**
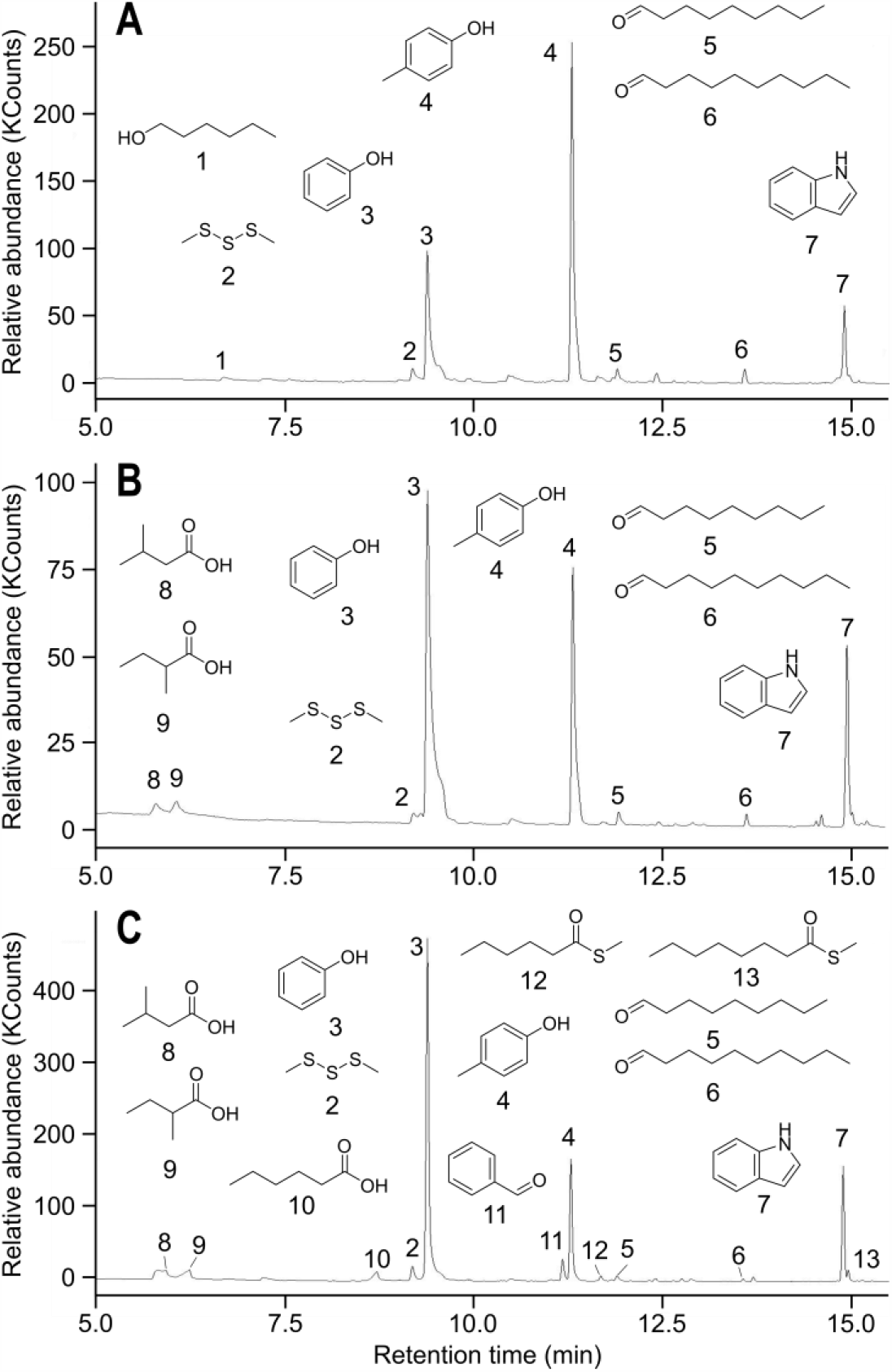
Composition of headspace odorants. Headspace odorants from (A) Brie cheese infusion, (B) Raclette cheese infusion, and (C) Limburger cheese infusion. Compounds were identified as 1-hexanol (1); dimethyl trisulfide (2); phenol (3); p-cresol (4); nonanal (5); decanal (6); indole (7); 3-methylbutyric acid (8); 2-methylbutyric acid (9); hexanoic acid (10); benzaldehyde (11); S-methyl hexanethioate (12); and S-methyl octanethioate (13).

#### Effect of cheese infusions as oviposition stimulants for mosquitoes (field experiment)

To determine the effect of a cheese infusion as oviposition stimulant for wild mosquitoes in field settings, we prepared a Brie cheese infusion (treatment), and a bluegrass, *Poa* sp., infusion (positive control), by allowing outdoor fermentation of either cheese or grass (50 g each) in 5 L of water for 5 days. In a paired-choice experiment (Figure 4A), the plastic tub filled with the cheese infusion received more egg rafts than the tub filled with the bluegrass infusion (n = 15, z = 2.9, p = 0.004; Figure 4B), but the proportional representation of egg rafts from female *Cx. pipiens* (35-38%) and female *Culiseta incidens* (62-65%) was similar for each type of infusion. Interestingly, we also captured (by drowning in tubs) males and females of *D. suzukii*, significantly more in the Brie cheese infusion than in the bluegrass infusion (n = 5, z = 5.00, p < 0.0001; Figure 4C). To consolidate these unexpected results, we placed paired tubs containing the Brie cheese infusion or a water control (Figure 4A) next to Himalayan blackberry, *Rubus armeniacus*, a *D. suzukii* host plant [67], and found that tubs with the Brie cheese infusion captured significantly more *D. suzukii* (n = 4, z = 6.25, p < 0.0001; Figure 4D). By-captures of other dipterans were not identified.

**Fig. 4.**
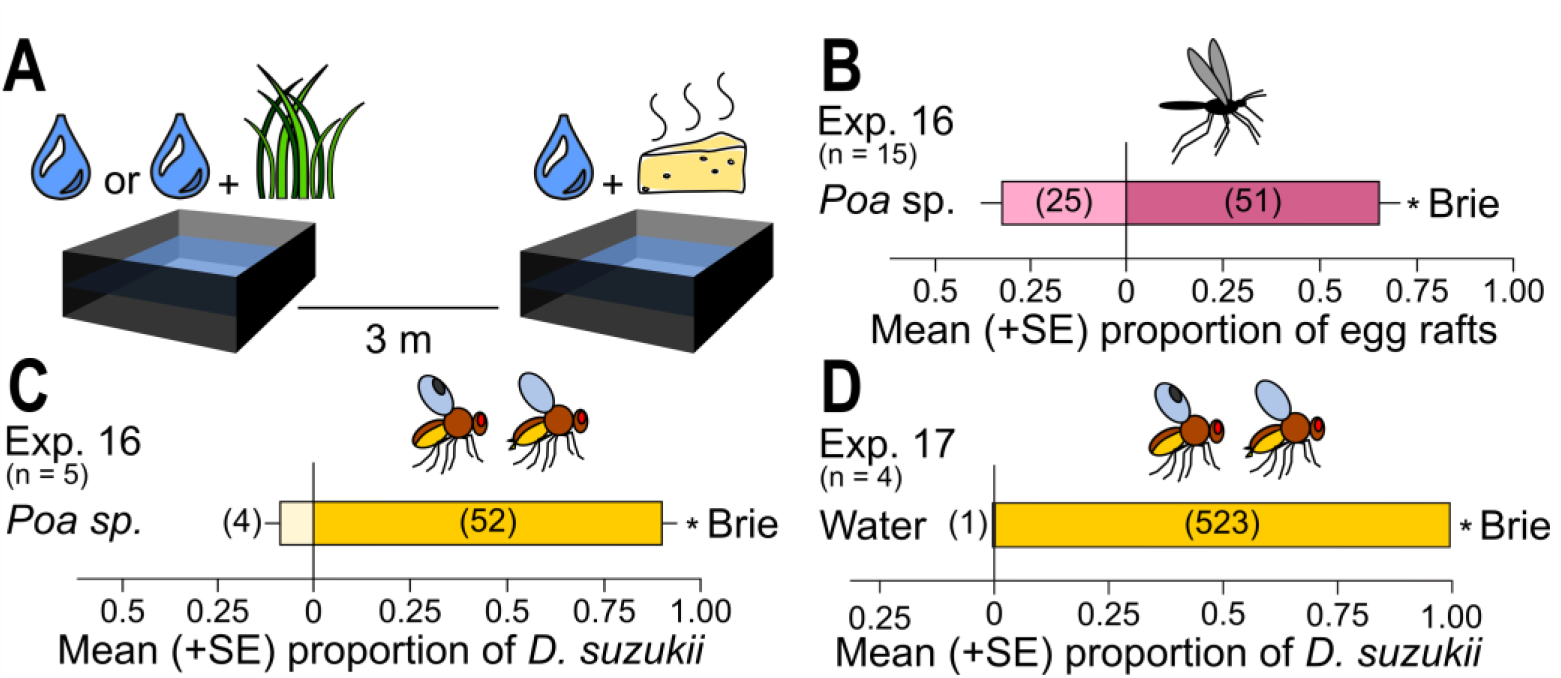
Effect of cheese infusion, or bluegrass infusion, on oviposition by mosquitoes and spotted wing Drosophila in field settings. **(A)** Graphical illustration of the experimental design applied in **B-D. (B, C)** Effect of Brie cheese infusion, or *Poa* sp. (bluegrass) infusion, on oviposition by *Culiseta incidens* and *Culex pipiens* **(B)**, and captures (by drowning) of *Drosophila suzukii* **(C). (D)** Effect of Brie cheese infusion or water on captures (by drowning) *of D. suzukii*. An asterisk indicates a significant preference (p < 0.05) for the specific test stimulus (binary logistic regression analysis with logit link function). Numbers within brackets indicate the total number of mosquito egg rafts placed, or *D. suzukii* captured, in response to the test stimulus.

#### Effect of microbe composition in home-made cheese infusions on oviposition choices made by mosquitoes (laboratory experiments)

To determine the effect of microbe composition in home-made cheese infusions on oviposition choices by mosquitoes, we prepared three types of cheese by adding different microbe starter cultures to skim milk powder and water, as follows: (i) mesophilic aromatic type B cheese: *Lactococcus lactis* ssp. *cremoris, L. lactis* ssp. *diacetylactis, L. lactis* ssp. *lactis*, and *Leuconostoc mesenteroides* ssp. *cremoris*; (ii) thermophilic type C cheese: *Lactobacillus helveticus, Streptococcus thermophilus*, and *Propionibacterium freudenreichii* ssp. *Shermanii*; and (iii) mesophilic type II cheese: *Lactococcus lactis* ssp. *Cremoris*. We then prepared three types of infusions by adding one of these three home-made cheeses to 180 mL of water and allowing indoor fermentation for 5 days. Given a choice between a cheese infusion and a water control, female *Cx. pipiens* placed more egg rafts on mesophilic aromatic type B infusions than on water controls (n = 8, z = 3.4, p <0.001), as many egg rafts on mesophilic type II cheese infusions and water controls (n = 8, z = 0.4, p < 0.69), and fewer egg rafts on mesophilic type II cheese infusions than on water controls (n = 8, z = − 5.2, p < 0.0001; Figure 5).

**Fig. 5.**
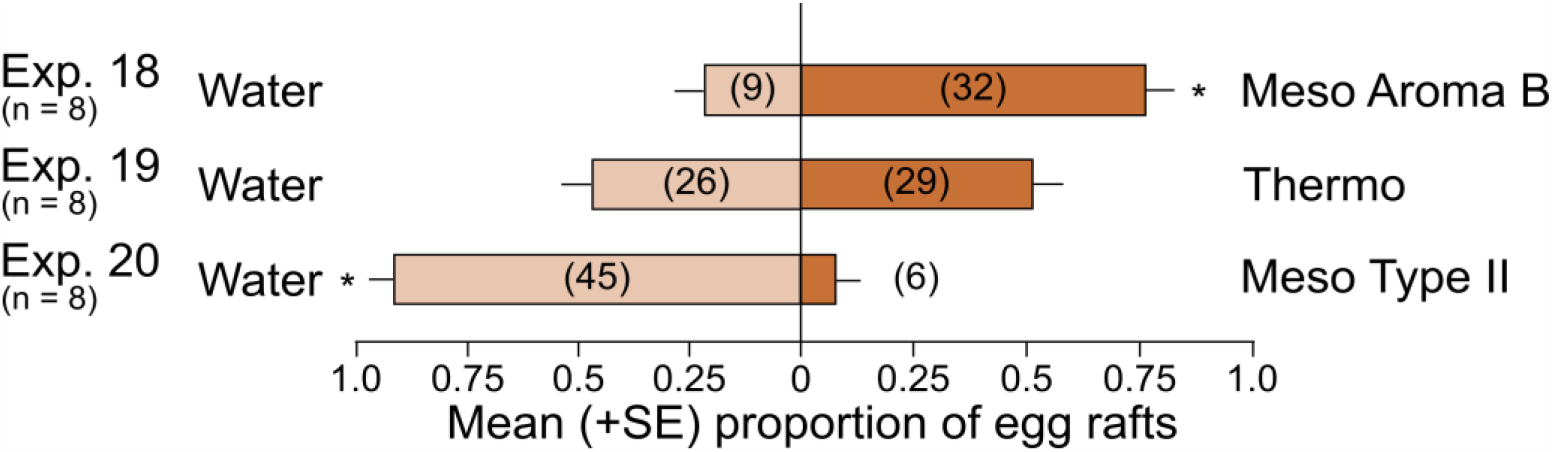
Effect of microbe composition in home-made cheese infusions on oviposition choices by mosquitoes. Selection of oviposition sites by groups of 10 female *Culex pipiens* each (8-15 days old; 4-5 days post blood-feeding) in response to contrasting microbe compositions represented in three cheese varieties [mesophilic aromatic type B (“Meso Aroma B), thermophilic type C (“Thermo”), mesophilic type II (Meso Type II)] used in cheese infusion. An asterisk indicates a significant preference (p < 0.05) for the specific test stimulus (binary logistic regression analysis with logit link function). Numbers within brackets indicate the total number of egg rafts placed in response to the test stimulus.

#### Effect of infusion type and nutrient provisioning on survivorship of mosquito larvae

To determine the effect of infusion type and nutrient provisioning on survivorship of mosquito larvae (10 first instars per cup), we (a) prepared infusions of home-made mesophilic aromatic type B cheese (20 g in 180 mL of distilled water) and of bluegrass (5 g in 195 mL of distilled water) allowing indoor fermentation for 5 days; (b) tested both infusion types in combination; (c) supplemented each infusion type, and both types in combination, with fish food every other day; and (d) used fish food alone as a positive control. There was a significant effect of treatment on larval survival (χ2 = 187.89, p < 0.0001; Figure 6). Fewer larvae survived in cheese infusions with fish-food supplement than in (i) cheese infusions alone (z = 2.68, p < 0.001), (ii) combined infusions of cheese and bluegrass with fish-food supplement (z = 2.58, p = 0.01), (iii) water with fish-food (z = 5.04, p < 0.0001), (iv) bluegrass infusions (z = 4.93, p < 0.0001), and (v) bluegrass infusions with fish-food supplement (z = 4.46, p < 0.0001). Similar number of larvae survived in cheese infusions, and in combined infusions of cheese and bluegrass with fish-food supplement (z = 0.38, P = 0.70); however, compared to each of these two treatments, fewer larvae survived in bluegrass infusions (z = 8.67, p < 0.0001; z = 8.68, p < 0.0001), bluegrass infusions with fish-food supplement (z = 7.64, p < 0.0001; z = 7.66, p < 0.0001), and in water with fish-food (z = 8.77, p < 0.0001; z = 8.79, p < 0.0001). Survivorship of larvae was similar in bluegrass infusions and in water with fish-food (z = 0.42, p = 0.67), and in bluegrass infusions and bluegrass infusions with fish-food supplement (z = 1.95, p = 0.052), but differed between bluegrass infusions with fish-food supplement and water with fish-food (z = 2.33, p = 0.02).

**Fig. 6.**
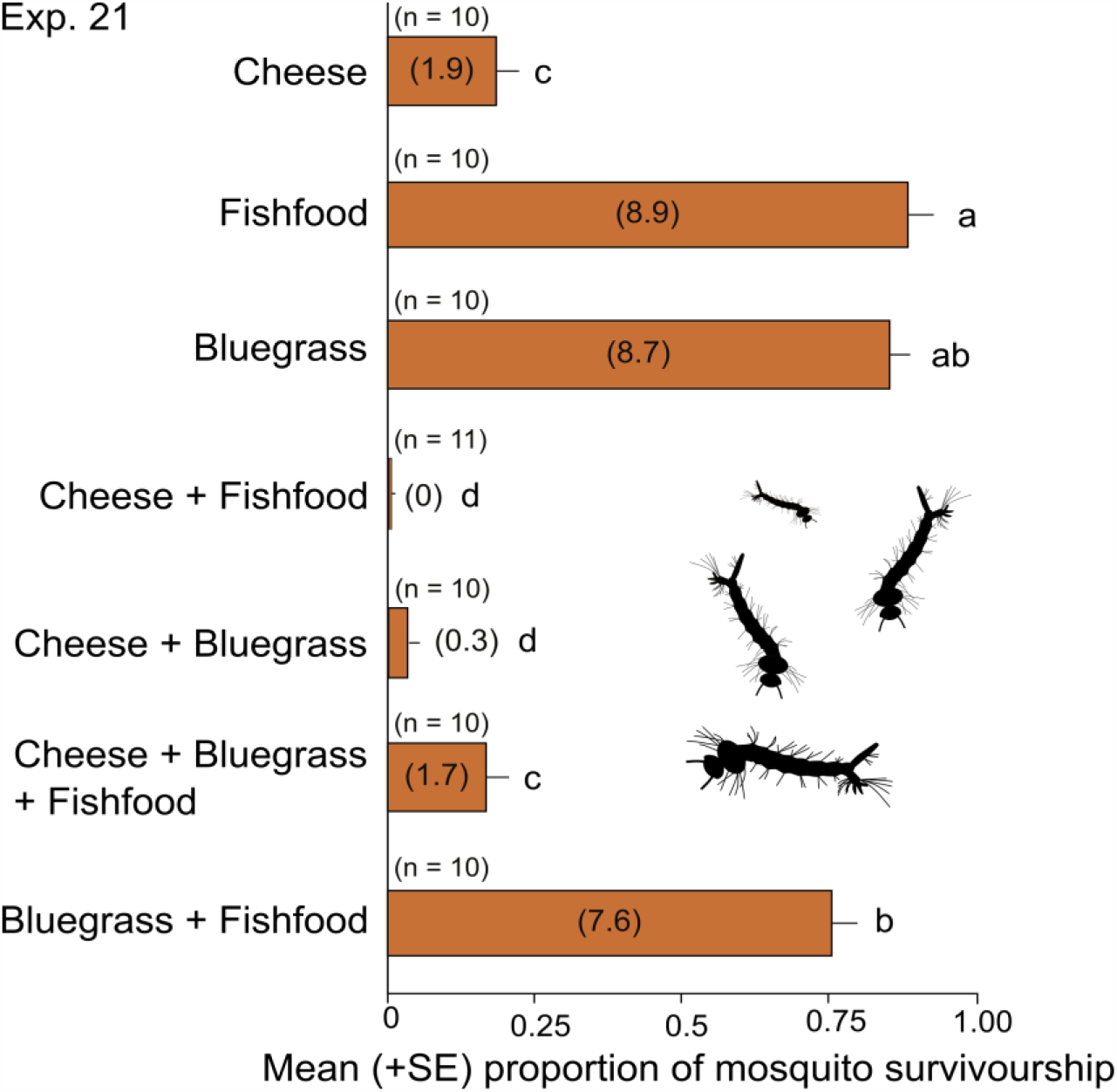
Larval habitat characteristics determine survivorship of mosquito larvae. Effects of infusion type (mesophilic aromatic type B cheese, bluegrass (*Poa* sp.)), and fish food supplement on the proportion of *Culex pipiens* larvae reaching adulthood. Different letters on bars indicate differences in the mean proportion of mosquitoes reaching adulthood (type 3 analysis of effects with Tukey’s HSD, p < 0.05); numbers within brackets indicate the mean number of larvae surviving to adults.

## Discussion

Our study (i) reveals the mechanisms underlying the attractiveness of cheese or cheese infusions to mosquitoes and SWDs seeking hosts or oviposition sites, (ii) explores cheese infusions as ecological traps for mosquitoes and SWDs, and (iii) promotes the concept of using affordable home-made cheese or cheese infusions for mosquito and SWD control.

Several species of mosquitoes, including *Ae. aegypti*, have been shown to be attracted to Limburger cheese, or its odor, in laboratory but not in field settings [50, 68]. We expand on these previous findings by showing attraction of host-seeking female *Ae. aegypti* and *Cx. pipiens* to various types of cheese (Raclette, Pecorino, Gruyere, Limburger, Brie) in laboratory experiments (Figure 1B), and attraction of *Cx. pipiens* and *Cs. incidens* to Brie in a field study (Figure 1D).

The attractiveness of cheese to mosquitoes is linked to the procedure and the ingredients for making cheese. Cheese production commonly involves select microbes that metabolize milk constituents and, in the process, generate lactic acid, fatty acids, amines, and even CO2 [58, 59, 66], all of which are important mosquito attractants [65, 69–71]. Lactic acid, e.g., is a major metabolite produced by members of the Gram-positive bacteria *Lactobacillales* which are commonly used in cheese-making [58, 59] and are part of the human skin microbiome [72]. The commonality of metabolites produced by these bacteria processing milk-based or skin-derived nutrients provide an ecological explanation for the attractiveness of cheese to host-seeking mosquitoes. As another dipteran, *D. suzukii* also exploits microbe-emitted odorants to locate fruit both as a food source and as an oviposition site [73]. Indeed, synthetic replicas of these microbe odorants are used as trap lures for *D. suzukii* [74, 75].

The ‘mother knows best’ principle (also known as the preference-performance hypothesis) applies to oviposition choices by female mosquitoes [76] and *D. suzukii* [77]. This principle was originally conceived for insect herbivores, and states that female insects select and oviposit on plants that maximise the survival and performance of their larvae, particularly when they cannot relocate and depend on their mother’s choice of host-plant [78, 79]. Female mosquitoes are under the same type of selection pressure. They commonly oviposit, and larvae develop, in discrete aquatic habitats such as small pools of water from which the larvae cannot relocate. Thus, selection of suitable larval habitats, and active avoidance of toxic habitats, by a gravid female mosquito is of paramount importance for the development and survival of her offspring [80]. It is perplexing then that female *Cx. pipiens* in our study oviposited in water pools with cheese infusions, which are lethal to their larval offspring. For mosquitoes, cheese infusions may be some kind of an ecological trap, representing a mismatch between a habitat’s perceived attractiveness and its actual quality, resulting in a population sink [39–41]. For example, aquatic oviposition sites containing foliar material of common blackberry, *Rubus allegheniensis*, represent an ecological trap for *Cx. pipiens* females, prompting them to oviposit but later killing their larval offspring [41]. Analogously, cheese infusions may signal suitable habitat at the time of oviposition but then become deleterious for larval development, possibly due to the built up of microbial waste or metabolites.

Selection of oviposition sites differed between mosquito taxa, reflecting the type of water body typically inhabited by larvae. Females of *Cx. pipiens* placed more egg rafts on water with Brie or Raclette cheese infusion than on plain water or water with Limburger cheese infusion (Figure 2A). Conversely, females of *Ae. aegypti* preferentially oviposited in plain water (Figure 2B), apparently avoiding any type of cheese infusion. Interestingly, both *Cx. pipiens* and *Cs. incidens* selected Brie cheese infusion over bluegrass infusion (Figure 4B), a known oviposition attractant for *Culex* spp. [35]. The oviposition choices by these mosquitoes likely reflect their habitat preference for larval development. Larvae of *Cx. pipiens* and *Cs. incidens* are often found in foul or polluted water with heavy loads of organic material, such as sewage lagoons, latrine pits, pools of agricultural runoff, or polluted storm drains [82–85], whereas larvae of *Ae. aegypti* often develop in tree holes or containers holding relatively clean water [82]. The unexpected attraction and capture (by drowning) of wild *D. suzukii* in cheese infusions (Figure 4C,D) may be attributed to microbial odorants resembling those emanating from microbe-infested fruit [22]. It seems possible to customize oviposition sites by engineering cheese infusions with select microbes and growth media for optimal attraction of mosquitoes or other insects (e.g., *D. suzukii*) that use similar odor cues to locate oviposition resources.

Gravid female mosquitoes are guided to oviposition sites by semiochemicals that are microbe-derived or originate from decaying organic material [36, 80], possibly indicative of habitat quality [86]. The odor profile of cheese infusions differed in accordance with the microbes and growth media involved (Figure 3), and apparently affected oviposition choices by gravid female *Cx. pipiens*. The specific headspace odorants engendered by a cheese infusion reflect interactions between the microbes selected for the infusion [87], the nutrients available to them [31, 88], and specific biotic and abiotic conditions [87]. Several odorants in the headspace of cheese infusions (Figure 3) including indole, phenol, p-cresol, and dimethyl trisulfide are known to dose-dependently function as oviposition attractants or stimulants for *Culex* mosquitoes [89, 90]. Indole is a major constituent of incubated human sweat [91], a quorum-sensing signal of bacteria [92], and an interkingdom signal facilitating communication between microbes and other organisms [92]. Dimethyl trisulfide, produced by the bacteria Lactococcus lactis and *Leuconostoc mesenteroides* breaking down carrion [99], attracts gravid callophorid flies to carrion oviposition sites [93]. These two microbes are commonly co-opted to make Brie and other soft cheeses [59].

Semiochemical baits, used alone or as part of an “attract and kill” tactic [94], often comprise blends of multiple semiochemicals that are formulated at the same ratio as they originated from the natural resource. This ratio, however, is commonly distorted when synthetic semiochemicals passively emanate from commercial dispensers [95]. If foraging insects are guided by semiochemical blends with “proper” composition and ratio of blend constituents to locate resources, such as human hosts, then distorted blends of synthetic semiochemicals may misguide foraging insects [96, 97] and fail to achieve the intended control effect in pest management programs [96, 97]. The deployment of metabolically-active, semiochemical-emitting microbes offers an appealing alternative to synthetic semiochemical lures for the control of host-seeking mosquitoes. This tactic could be optimized by carefully “designing” the nutritional composition of media for microbial growth, selecting key microbes for media inoculation, and by extending their metabolic activity through slow-release formulations of essential nutrients. If some microbial metabolites were to be both attractive and toxic to mosquitoes seeking hosts and/or oviposition sites (see above), then these microbial metabolites could be used for both the attract and the kill function of “attract and kill” tactics.

Insect pest burdens often fall disproportionately on underdeveloped regions [98] which have limited resources for pest management. Home-made cheese infusions are relative affordable and have the potential for use in integrated programs aimed at control of mosquitoes and other pest insects. The use of customizable and non-conventional nutritional media, such as cheese, as microbe-based ecological traps presents a promising concept which exploits insect ecology for insect control.

## Materials and Methods

### Rearing of mosquitoes

We reared mosquitoes under a photoperiod of 14L:10D at 23-26 °C and 40-60% RH. We maintained mixed groups of males and females in mesh cages (30 × 30 × 46 cm high) provisioned ad libitum with a 10-% sucrose solution. Once per week, DP fed female mosquitoes on his arm. For oviposition, gravid Cx. pipiens females were given access to water in a circular glass dish (10 cm diameter × 5 cm high), and gravid Ae. aegypti females were given access to a 354-mL cup (Solo Cup Company, Lake Forest, IL 60045, USA) with paper towel lining (Kruger Inc., Montréal, QC H3S 1G5, Canada). We transferred Cx. pipiens egg rafts to water-filled trays (45 × 25 × 7 cm high), and paper towel strips with Ae. aegypti eggs to circular glass dishes (10 cm diameter × 5 cm high) containing water and brewers yeast (U.S. Biological Life Sciences, Salem, MA 01970, USA), and 2-4 days later transferred the dish contents to water-filled trays (45 × 25 × 7 cm high). We provisioned larvae with NutriFin Basix tropical fish food (Rolf C. Hagen Inc., Baie-D’Urfe, QC H9X 0A2, Canada), and transferred pupae via a 7-mL plastic pipette (VWR International, Radnor, PA 19087, USA) to water-containing 354-mL Solo cups covered with a mesh lid. We collected eclosed adults via aspirator and placed them in similar cups, along with a cotton ball soaked in a 10-% sucrose solution.

### Blood feeding for oviposition experiments

We starved one-week-old female mosquitoes for 24 h prior to blood feeding. To this end, we pipetted defibrillated sheep blood into a 50-mL centrifuge tube (VWR International), and then covered its mouth with a thin, stretched layer of Parafilm M (Bemis Company Inc., Neenah, WI 54956, USA) [99]. The tube was heated to 37 °C in a water bath and placed mouth-down on top of a mesh-covered cup (Solo Cup Company), allowing the mosquitoes to blood-feed through the mesh for 30 min. We then aspirated engorged mosquitoes into wooden mesh cages (30 × 30 × 46 cm high) provisioning them with a 10-% sucrose solution ad libitum.

### Cheese as an attractant to host-seeking mosquitoes in laboratory settings

Using a paired-trap experimental design (Figure 1A), we ran behavioural bioassays in mesh cages (77 × 78 × 104 cm) wrapped with black cloth except for the top to allow illumination from ambient fluorescent light. We kept bioassay conditions at 23-26 °C, 40-60% RH, and a photoperiod of 14L:10D. For each 24-h bioassay, we released 50 virgin, 2-to 10-day-old, 24-h sugar-deprived females of Ae. aegypti or Cx. pipiens from a Solo cup into a cage. We randomly assigned the treatment and the control stimulus to paired adhesive-coated (The Tanglefoot Comp., Grand Rapids, MI 49504, USA), custom-made delta traps (9 cm × 15 cm) placed on stands spaced 30 cm apart inside the cage. Wearing latex gloves (Microflex Corp., Reno, NV 89523, USA), we fitted each trap with a Petri dish (100 × 15 mm) (VWR International), one of which contained 20 g of a cheese variety [Brie (Castello Cheese Inc., Concord, ON L4K 2G9, Canada), Mozzarella (Western Family Foods, Tigard, OR 97223, USA), Raclette (Emmi Cheese, 6005 Luzern, Switzerland), Gruyère (Emmi Cheese, 6005 Luzern, Switzerland), Parmesan (Maison Riviera, Varennes, QC J3X 1P7, Canada), Pecorino (Saputo Inc., Montreal, QC H1P 1×8, Canada), Cheddar (Black Diamond Cheese Ltd., Belleville, ON K8N 5A1, Canada), Limburger (Agropur Cooperative, Saint-Hubert, QC J3Z 1G5, Canada)] as the treatment stimulus. Both the treatment and the control dish received a wet cotton ball (Fisher Scientific, Pittsburgh, PA 02451, USA) to ensure equal humidity between the two test stimuli.

### Cheese as an attractant to host-seeking mosquitoes in field settings

For each replicate (n = 15), we placed two BG sentinel traps (Biogents AG, Regensburg, Germany) 10-m apart on a lawn on the Burnaby campus of Simon Fraser University (SFU), 3 m away from a patch of vegetation. We baited each trap with carbon dioxide (CO2) emanating from 1 kg of dry ice and added 20 g of Brie (see above), cut into four pieces, to the treatment trap. We commenced experimental replicates about 30 min before sunset and terminated them about 90 min later.

### Preparation of oviposition stimulants

We prepared infusions of store-bought cheese (treatments) or bluegrass, Poa sp., (positive control) to test their effect as oviposition stimulants for mosquitoes. To this end, we added 50 g of a commercial cheese [Brie (Castello Cheese Inc.); Raclette (Emmi Cheese); Limburger: (Agropur Cooperative)], or 50 g of blue grass collected on the Burnaby campus of SFU, to 5 L of distilled water, and allowed fermentation to proceed in an outdoor shelter for 5 days. The negative control consisted of 5 L of distilled water steeping under the same conditions for 5 days.

We also prepared three types of infusions with home-made cheese: mesophilic aromatic type B, mesophilic type II, and thermophilic type C (all ingredients from Biena Inc., St. Hyacinthe, QC J2S 1L4, Canada). We made mesophilic aroma type B cheese by mixing 925 mL of water, 300 mg of skim milk powder (Compliments skim milk powder, Sobeys Inc., Stellarton, NS B0K 1S0, Canada), and 1 g of mesophilic aromatic type B starter culture (Lactococcus lactis ssp. cremoris, L. lactis ssp. diacetylactis, L. lactis ssp. lactis, Leuconostoc mesenteroides ssp. cremoris); we made mesophilic type II cheese by mixing 925 mL of water, 300 mL of skim milk powder, and 1 g of mesophilic type II starter culture (Lactococcus lactis ssp. cremoris) (Biena Inc.); we made thermophilic type C cheese by mixing 925 mL of water, 300 mL of skim milk powder, 1 g of thermophilic starter culture type C (Lactobacillus helveticus and Streptococcus thermophilus) (Biena Inc.), and 1 g Propionibacterium 50 (Propionibacterium freudenreichii ssp. shermanii) (Biena Inc.). We prepared infusions of these home-made cheeses by adding 20 g of home-made cheese to 180 mL of distilled water, and we prepared a grass infusion by adding 5 g of bluegrass to 195 mL of distilled water. We allowed indoor fermentation of these infusions for 5 days at 23-26 °C, 40-60% RH, and a photoperiod of 14L:10D.

### Infusions of store-bought cheese as oviposition stimulants for mosquitoes in laboratory experiments

We ran all bioassays at 23-26 °C, 40-60% RH, and a photoperiod of 14L:10D. For each replicate, we aspirated 20 females of Ae. aegypti or Cx. pipiens, 8-15 days old and 4-5 days post blood feeding, into an aluminum mesh cage (61 × 61 × 61 cm) (BioQuip Products Inc., Compton, CA 90220, USA) fitted with paired cups (Solo Cup Company) containing either 200 mL of a cheese infusion or water (control). After 48 h, we removed the cups and counted the number of Cx. pipiens egg rafts, or the number of Ae. aegypti eggs, placed in each cup. In each of experiments 12 (Cx. pipiens) and 13 (Ae. aegypti), we tested four oviposition stimulants: (1) Brie cheese infusion, (2) Raclette cheese infusion, (3) Limburger cheese infusion, and (4) water. For each replicate, we placed a cup with 200 mL of a test stimulus in each corner of the mesh cage, with 20 cm spacing between cups. In experiment 14 (Cx. pipiens), we tested the effect of two cheese infusions (Brie vs Raclette) head-to-head by placing the two cups with a test stimulus 20 cm apart on opposite sides of a mesh cage.

### Dynamic collection of headspace odorants from cheese infusions and their effect on oviposition by mosquitoes

We placed 200 mL of a Brie cheese infusion into a Pyrex® glass chamber (34 cm high × 12.5 cm wide), and with a mechanical pump pulled charcoal-filtered air at a flow of 1 L min-1 for 24-72 h through the chamber and through a glass column (6 mm outer diameter × 150 mm) containing 200 mg of Porapak-Q™ adsorbent [100]. We desorbed cheese infusion odorants captured on Porapak-Q with 2 mL of pentane and ether (1:1), and for a test (treatment) stimulus diluted 100-µL aliquots of this headspace odorant extract (HOE) in 900 µL of pentane and ether (1:1 mix). To test the effect of this stimulus on selection of oviposition sites by mosquitoes (Exp. 15), we pipetted the dilution into a 4-mL glass vial with a 2-mm hole in its lid, replacing the dilution with pentane/ether (1:1, 1 mL) as the control stimulus. We randomly assigned the treatment and the control stimulus to paired cups (see above) filled with 200 mL of water and placed 20 cm apart on opposite sides of a mesh cage. We then introduced 10 females of Cx. pipiens (8-15 days old, 4-5 days post blood feeding) and allowed 48 h for oviposition.

### Identification of odorants in headspace odorant extracts (HOEs)

We analyzed 2-µl aliquots of HOE by gas chromatography-mass spectrometry (GC-MS), operating a Saturn 2000 Ion Trap GC-MS fitted with a DB-5 GC-MS column (30 m × 0.25 mm i.d.; Agilent Technologies Inc., Santa Clara, CA 95051, USA) in full-scan electron impact mode. We used a flow of helium (35 cm s-1) as the carrier gas with the following temperature program: 50 °C (5 min), 10 °C min-1 to 280 °C (held for 10 min). The temperature of both the injector port and ion trap was 250 °C. To identify headspace odorants, we compared their retention indices [101] and mass spectra with those of authentic standards for 1-hexanol, dimethyl trisulfide, phenol, p-cresol, nonanal, decanal, indole, 3-methylbutyric acid, 2-methylbutyic acid, hexanoic acid and benzaldehyde (all from Sigma-Aldrich, St. Louis, MO 63103 MO, USA). S-Methyl hexanethioate and S-methyl octanethioate were synthesized from methyl hexanoate and methyl octanoate (Sigma-Aldrich), respectively, following established methods [102].

### Infusions of Brie cheese, or bluegrass, as oviposition stimulants for mosquitoes in field settings

To compare the effect of cheese (Brie) or bluegrass infusions (see above for preparation) on the selection of oviposition sites by wild mosquitoes (Exp. 16, n = 15, July 2018), we placed paired black plastic tubs (38 × 50 × 13 cm high) (Biogents AG, Regensburg, Germany) with 3-m inter-tub spacing on the Burnaby campus of Simon Fraser University (SFU, BC, Canada), and baited each tub with 5 L of the brie or bluegrass infusion. After 48 h, we counted the number of egg rafts present and transferred each raft into a separate cup (Solo Cup Company) to allow larval development and identification of emergent adults to species. Partway through experiment 16, we noticed the capture (by drowning) of spotted-wing drosophila, Drosophila suzukii, in Brie infusion tubs.

### Effect of Brie cheese infusion on attraction of D. suzukii in field settings

To test the effect of Brie cheese infusion on attraction and capture of D. suzukii (Exp. 17, n = 4, July-August 2018), we placed paired black plastic tubs (see above) within 3 m of Himalayan blackberry patches on the SFU Burnaby campus, and baited each tub with 5 L of the Brie cheese infusion or distilled water. After 48 h, we counted the number of D. suzukii drowned in each tub.

### Effect of microbe composition in home-made cheese infusions on oviposition choices by mosquitoes

To test the effect of microbe composition in home-made cheese infusions on oviposition choices by mosquitoes, we prepared separate infusions from mesophilic aromatic type B, mesophilic type II, and thermophilic starter culture type C (see preparation of oviposition stimulants). For each experimental replicate in experiments 18-20, we inserted paired oviposition cups (Solo Cup Company) into an aluminum mesh cage (61 × 61 × 61 cm) (BioQuip Products Inc.), each cup containing 200 mL of a cheese infusion or a water control. We then introduced 20 gravid Cx. pipiens females (5 days post blood feeding) and allowed them to select oviposition sites during 48 h, maintaining ambient conditions at 23-26 °C, 40-60% RH, and a photoperiod of 14L:10D. Thereafter, we removed oviposition cups and counted the number of egg rafts they had received.

### Effect of infusion type and nutrient provisioning on survivorship of mosquito larvae

To test the effect of infusion type and nutrient provisioning on survivorship of mosquito larvae (Exp. 21, n = 10), for each replicate, we pipetted 10 first instar Cx. pipiens larvae into a 200-mL water reservoir covered with a mesh lid, testing 7 treatments: (1) mesophilic type B cheese infusion, (2) bluegrass infusion, (3) clean water with 0.1 mg of NutriFin Basix tropical fish-food (Rolf C. Hagen Inc.) added every other day, (4-5) cheese infusion in binary combination with either fish-food (4) or bluegrass infusion (5), (6) cheese infusion in ternary combination with bluegrass infusion and fish-food, and (7) bluegrass infusion in binary combination with fish-food. We skimmed the water surface daily to remove any microbial build-up, and recorded adult emergence. Ambient conditions during larval development were: 23-26 °C, 40-60% RH, and a photoperiod of 14L:10D.

### Statistical analyses

We used SAS statistical software version 9.4 (SAS Institute Inc., Cary, NC 27513, USA) for data analyses. We used a binary logistic regression model with a logit link function to compare mean proportions of responders between test stimuli. In addition, for experiments 12, 13, and 21 we used a type 3 analysis of effects to test for effect presence, and a Tukey’s HSD to determine pairwise differences between mean effects. We worked with back-transformed data to obtain means and confidence intervals.

## End Matter

### Author Contributions and Notes

Conceptualization, D.A.H.P. and M.A.; Methodology, D.A.H.P., R.G., and G.G.; Investigation, D.A.H.P., M.A., E.K., S.M., R.G.; Writing – Original Draft, D.A.H.P., M.A., R.G., and G.G., Funding Acquisition, G.G.; Resources, G.G. The authors declare no conflict of interest.

This article contains supporting information online. <if true>

## Acknowledgments

We thank Martin Duckhorn and Grace Peach for volunteer assistance in the project and Bernhard Senge for discussion.

## References

1. Devi, S. (2020). Locust swarms in east Africa could be “a catastrophe.” The Lancet 395, 547.

2. A lack of locust preparedness will cost lives (2020). Nature 579, 174–174.

3. Goulson, D., Nicholls, E., Botias, C., and Rotheray, E.L. (2015). Bee declines driven by combined stress from parasites, pesticides, and lack of flowers. Science 347, 1255957–1255957.

4. Wagner, D.L. (2020). Insect declines in the Anthropocene. Annu. Rev. Entomol. 65, 457–480.

5. Gippet, J.M., Liebhold, A.M., Fenn-Moltu, G., and Bertelsmeier, C. (2019). Human-mediated dispersal in insects. Curr. Opin. Insect Sci. 35, 96–102.

6. Bradshaw, C.J.A., Leroy, B., Bellard, C., Roiz, D., Albert, C., Fournier, A., Barbet-Massin, M., Salles, J.-M., Simard, F., and Courchamp, F. (2016). Massive yet grossly underestimated global costs of invasive insects. Nat. Commun. 7, 12986.

7. Gates, B. (2014). The Deadliest Animal in the World. gatesnotes.com. https://www.gatesnotes.com/Health/Most-Lethal-Animal-Mosquito-Week.

8. Hemingway, J., Ranson, H., Magill, A., Kolaczinski, J., Fornadel, C., Gimnig, J., Coetzee, M., Simard, F., Roch, D.K., Hinzoumbe, C.K., et al. (2016). Averting a malaria disaster: will insecticide resistance derail malaria control? The Lancet 387, 1785–1788.

9. Dang, K., Doggett, S.L., Veera Singham, G., and Lee, C.-Y. (2017). Insecticide resistance and resistance mechanisms in bed bugs, Cimex spp. (Hemiptera: Cimicidae). Parasit. Vectors 10, 318.

10. Potts, S.G., Imperatriz-Fonseca, V., Ngo, H.T., Aizen, M.A., Biesmeijer, J.C., Breeze, T.D., Dicks, L.V., Garibaldi, L.A., Hill, R., Settele, J., et al. (2016). Safeguarding pollinators and their values to human well-being. Nature 540, 220–229.

11. Pimentel, D., and Burgess, M. (2014). Environmental and Economic Costs of the Application of Pesticides Primarily in the United States. In Integrated Pest Management: Pesticide Problems, Vol. 3, D. Pimentel and R. Peshin, eds. (Springer Netherlands), pp. 47–71.

12. Blancke, S., Van Breusegem, F., De Jaeger, G., Braeckman, J., and Van Montagu, M. (2015). Fatal attraction: the intuitive appeal of GMO opposition. Trends Plant Sci. 20, 414–418.

13. Suckling, D.M., Stringer, L.D., Stephens, A.E., Woods, B., Williams, D.G., Baker, G., and El-Sayed, A.M. (2014). From integrated pest management to integrated pest eradication: Technologies and future needs. Pest Manag. Sci. 70, 179–189.

14. Ferguson, H.M., Dornhaus, A., Beeche, A., Borgemeister, C., Gottlieb, M., Mulla, M.S., Gimnig, J.E., Fish, D., and Killeen, G.F. (2010). Ecology: A prerequisite for malaria elimination and eradication. PLoS Med. 7, e1000303.

15. Huang, W., Wang, S., and Jacobs-Lorena, M. (2020). Use of microbiota to fight mosquito-borne disease. Front. Genet. 11, 196.

16. Wooding, M., Naudé, Y., Rohwer, E., and Bouwer, M. (2020). Controlling mosquitoes with semiochemicals: a review. Parasit. Vectors 13, 80.

17. Peterson, R.K.D., Higley, L.G., and Pedigo, L.P. (2018). Whatever happened to IPM? Am. Entomol. 64, 146–150.

18. Ali, M.P., Bari, M.N., Haque, S.S., Kabir, M.M.M., Afrin, S., Nowrin, F., Islam, M.S., and Landis, D.A. (2019). Establishing next-generation pest control services in rice fields: eco-agriculture. Sci. Rep. 9, 10180.

19. McFall-Ngai, M., Hadfield, M.G., Bosch, T.C.G., Carey, H.V., Domazet-Lošo, T., Douglas, A.E., Dubilier, N., Eberl, G., Fukami, T., Gilbert, S.F., et al. (2013). Animals in a bacterial world, a new imperative for the life sciences. Proc. Natl. Acad. Sci. 110, 3229– 3236.

20. Verhulst, N.O., Qiu, Y.T., Beijleveld, H., Maliepaard, C., Knights, D., Schulz, S., Berg-Lyons, D., Lauber, C.L., Verduijn, W., Haasnoot, G.W., et al. (2011). Composition of human skin microbiota affects attractiveness to malaria mosquitoes. PLoS ONE 6, e28991.

21. Burkepile, D.E., Parker, J.D., Woodson, C.B., Mills, H.J., Kubanek, J., Sobecky, P.A., and Hay, M.E. (2006). Chemically mediated competition between microbes and animals: Microbes as consumers in food webs. Ecology 87, 2821–2831.

22. Hamby, K.A., and Becher, P.G. (2016). Current knowledge of interactions between Drosophila suzukii and microbes, and their potential utility for pest management. J. Pest Sci. 89, 621–630.

23. Dethlefsen, L., McFall-Ngai, M., and Relman, D.A. (2007). An ecological and evolutionary perspective on human–microbe mutualism and disease. Nature 449, 811–818.

24. Werren, J.H., Baldo, L., and Clark, M.E. (2008). Wolbachia: Master manipulators of invertebrate biology. Nat. Rev. Microbiol. 6, 741– 751.

25. Misof, B., Liu, S., Meusemann, K., Peters, R.S., Donath, A., Mayer, C., Frandsen, P.B., Ware, J., Flouri, T., Beutel, R.G., et al. (2014). Phylogenomics resolves the timing and pattern of insect evolution. Science 346, 763.

26. Davis, T.S., Crippen, T.L., Hofstetter, R.W., and Tomberlin, J.K. (2013). Microbial volatile emissions as insect semiochemicals. J. Chem. Ecol. 39, 840–859.

27. Oliver, K.M., and Martinez, A.J. (2014). How resident microbes modulate ecologically-important traits of insects. Curr. Opin. Insect Sci. 4, 1–7.

28. Douglas, A.E. (2015). Multiorganismal insects: Diversity and function of resident microorganisms. Annu. Rev. Entomol. 60, 17–34.

29. Brysch-Herzberg, M. (2004). Ecology of yeasts in plant–bumblebee mutualism in Central Europe. FEMS Microbiol. Ecol. 50, 87–100.

30. Mueller, U.G., Gerardo, N.M., Aanen, D.K., Six, D.L., and Schultz, T.R. (2005). The evolution of agriculture in insects. Annu. Rev. Ecol. Evol. Syst. 36, 563–595.

31. Peach, D.A.H., Carroll, C., Meraj, S., Gomes, S., Galloway, E., Balcita, A., Coatsworth, H., Young, N., Uriel, Y., Gries, R., et al. (2020). Nectar-dwelling microbes of common tansy are attractive to its mosquito pollinator, Culex pipiens L. bioRxiv. doi:10.1101/2020.04.03.024380

32. Leroy, P.D., Sabri, A., Verheggen, F.J., Francis, F., Thonart, P., and Haubruge, E. (2011). The semiochemically mediated interactions between bacteria and insects. Chemoecology 21, 113–122.

33. Uriel, Y., Gries, R., Tu, L., Carroll, C., Zhai, H., Moore, M., and Gries, G. (2020). The fly factor phenomenon is mediated by inter-kingdom signalling between bacterial symbionts and their blow fly hosts. Insect Sci. 27, 256–265.

34. Babcock, T., Gries, R., Borden, J., Palmero, L., Mattiacci, A., Masciocchi, M., Corley, J., and Gries, G. (2017). Brewer’s yeast, Saccharomyces cerevisiae, enhances attraction of two invasive yellowjackets (Hymenoptera: Vespidae) to dried fruit and fruit powder. J. Insect Sci. 17, 91.

35. Takken, W., and Verhulst, N.O. (2017). Chemical signaling in mosquito–host interactions: the role of human skin microbiota. Curr. Opin. Insect Sci. 20, 68–74.

36. Ponnusamy, L., Xu, N., Nojima, S., Wesson, D.M., Schal, C., and Apperson, C.S. (2008). Identification of bacteria and bacteria-associated chemical cues that mediate oviposition site preferences by Aedes aegypti. PNAS 105, 9262–9267.

37. Leroy, P.D., Sabri, A., Heuskin, S., Thonart, P., Lognay, G., Verheggen, F.J., Francis, F., Brostaux, Y., Felton, G.W., and Haubruge, E. (2011). Microorganisms from aphid honeydew attract and enhance the efficacy of natural enemies. Nat. Commun. 2, 348.

38. Peach, D.A.H., Gries, R., Young, N., Lakes, R., Galloway, E., Alamsetti, S.K., Ko, E., Ly, A., and Gries, G. (2019). Attraction of female Aedes aegypti (L.) to aphid honeydew. Insects 10, 43.

39. Gates, J.E., and Gysel, L.W. (1978). Avian nest dispersion and fledging success in field-forest ecotones. Ecology 59, 871–883.

40. Weldon, A.J., and Haddad, N.M. (2005). The effects of patch shape on indigo buntings: Evidence for an ecological trap. Ecology 86, 1422–1431.

41. Gardner, A.M., Muturi, E.J., and Allan, B.F. (2018). Discovery and exploitation of a natural ecological trap for a mosquito disease vector. Proc. R. Soc. B Biol. Sci. 285, 20181962.

42. Matthews, B.J., Younger, M.A., and Vosshall, L.B. (2019). The ion channel ppk301 controls freshwater egg-laying in the mosquito Aedes aegypti. eLife 8, e43963.

43. Walton, W.E., Van Dam, A.R., and Popko, D.A. (2009). Ovipositional responses of two Culex (Diptera: Culicidae) species to larvivorous fish. J. Med. Entomol. 46, 1338–1343.

44. Hazard, E., Mayer, M., and Savage, K. (1967). Attraction and oviposition stimulation of gravid female mosquitoes by bacteria isolated from hay infusions. Mosq. News 27, 133–136.

45. Gemeno, S.A.R., Trexler, J.D., Apperson, C.S., Zurek, L., Schal, C., Kaufman, M., Walker, E., Watson, D.W., and Wallace, L. (2003). Role of bacteria in mediating the oviposition responses of Aedes albopictus (Diptera: Culicidae). J. Med 40, 841–848.

46. Verhulst, N.O., Beijleveld, H., Knols, B.G., Takken, W., Schraa, G., Bouwmeester, H.J., and Smallegange, R.C. (2009). Cultured skin microbiota attracts malaria mosquitoes. Malar. J. 8, 302.

47. Braks, M.A.H., and Takken, W. (1999). Incubated human sweat but not fresh sweat attracts the malaria mosquito Anopheles gambiae sensu stricto. J. Chem. Ecol. 25, 663–672.

48. De Jong, R., and Knols, B. (1995). Olfactory responses of host-seeking Anopheles gambiae s.s. Giles (Diptera: Culicidae). Acta Trop. 59, 333–335.

49. Knols, B., van Loon, J., Cork, A., and Takken, W. (1997). Behavioural and electrophysiological responses of female Anopheles gambiae to Limburger cheese volatiles. Bull. Entomol. Res. 87, 151–159.

50. Kline, D.L. (1998). Olfactory responses and field attraction of mosquitoes to volatiles from Limburger cheese and human foot odor. J. Vector Ecol. 23, 186–194.

51. De Jong, R., and Knols, B. (1995). Olfactory responses of host-seeking Anopheles gambiae s.s. Giles (Diptera: Culicidae).Acta Trop. 59, 333–335.

52. Candida, V., and Agatino, R. (2004). Piophila casei L. (Diptera: Piophilidae) monitoring in cheese ripening storehouses. Integr. Prot. Stored Prod. IOBC Bull. 27, 109–114.

53. Jones, B.R., Graham, P.P., and Kelly, R.F. (1971). Microorganisms as inducers of oviposition for the cheese skipper, Piophla casei (L) Diptera. J Milk Food Technol 34, 410–415.

54. Hegazi, E.M., El-Gayar, F.H., Rawash, I.A., and Ali, S.A. (1978). Factors affecting the bionomics of Piophila casei (L.). Z. für Angew. Entomol. 85, 327–335.

55. Knols, B., van Loon, J., Cork, A., Robinson, R., Adam, W., Meijerink, J., De Jong, R., and Takken, W. (1997). Behavioural and electrophysiological responses of the female malaria mosquito Anopheles gambiae (Diptera: Culicidae) to Limburger cheese volatiles. Bull. Entomol. Res. 87, 151–159.

56. Knols, B.G.J., and De Jong, R. (1996). Limburger cheese as an attractant for the malaria mosquito Anopheles gambiae s.s. Parasitol. Today 12, 159–161.

57. Sablé, S., and Cottenceau, G. (1999). Current knowledge of soft cheeses flavor and related compounds. J. Agric. Food Chem. 47, 4825–4836.

58. Cogan, T., and Beresford, T. (2002). Microbiology of hard cheese. In Dairy Microbiology Handbook: The Microbiology of Milk and Milk Products, R. Robinson, ed., pp. 555–560.

59. Farkye, N., and Vedamuthu, E. (2002). Microbiology of soft cheeses. In Dairy Microbiology Handbook: The Microbiology of Milk and Milk Products, R. Robinson, ed., pp. 479–514.

60. Smit, G., Smit, B.A., and Engels, W.J.M. (2005). Flavour formation by lactic acid bacteria and biochemical flavour profiling of cheese products. FEMS Microbiol. Rev. 29, 591–610.

61. Kongo, J. (2013). Lactic acid bacteria as starter-cultures for cheese processing: past, present and future developments. In Lactic Acid Bacteria − R & D for Food, Health and Livestock Purposes (Intech), pp. 1–21.

62. Fierer, N., Hamady, M., Lauber, C.L., and Knight, R. (2008). The influence of sex, handedness, and washing on the diversity of hand surface bacteria. Proc. Natl. Acad. Sci. 105, 17994–17999.

63. Endo, A., Irisawa, T., Futagawa-Endo, Y., Sonomoto, K., Itoh, K., Takano, K., Okada, S., and Dicks, L.M.T. (2011). Fructobacillus tropaeoli sp. nov., a fructophilic lactic acid bacterium isolated from a flower. Int. J. Syst. Evol. Microbiol. 61, 898–902.

64. Zaunmüller, T., Eichert, M., Richter, H., and Unden, G. (2006). Variations in the energy metabolism of biotechnologically relevant heterofermentative lactic acid bacteria during growth on sugars and organic acids. Appl. Microbiol. Biotechnol. 72, 421–429.

65. Acree, F., Turner, R.B., Gouck, H.K., Berzoa, M., and Smith, N. (1968). L-lactic acid: A mosquito attractant isolated from humans. Science 161, 1346–1347.

66. Tamime, A. (2002). Microbiology of starter cultures. In Dairy Microbiology Handbook, Third Edition, R. K. Robinson, ed. (John Wiley and Sons), pp. 261–366.

67. Lee, J.C., Dreves, A.J., Cave, A.M., Kawai, S., Isaacs, R., Miller, J.C., Van Timmeren, S., and Bruck, D.J. (2015). Infestation of wild and ornamental noncrop fruits by Drosophila suzukii (Diptera: Drosophilidae). Ann. Entomol. Soc. Am. 108, 117–129.

68. Murphy, M.W., Dunton, R.F., Perich, M.J., and Rowley, W. A. (2001). Attraction of Anopheles (Diptera: culicidae) to volatile chemicals in Western Kenya. J. Med. Entomol. 38, 242–244.

69. Eiras, A.E., and Jepson, P.C. (1994). Responses of female Aedes aegypti (Diptera: Culicidae) to host odours and convection currents using an olfactometer bioassay. Bull. Entomol. Res. 84, 207–211.

70. Bowen, M.F. (1991). The sensory physiology of host-seeking behavior in mosquitoes. Annu. Rev. Entomol. 36, 139–158.

71. Smallegange, R.C., Qiu, Y.T., van Loon, J.A., and Takken, W. (2005). Synergism between ammonia, lactic acid and carboxylic acids as kairomones in the host-seeking behaviour of the malaria mosquito Anopheles gambiae sensu stricto (Diptera: Culicidae). Chem. Senses 30, 145–152.

72. Grice, E.A., Kong, H.H., Conlan, S., Deming, C.B., Davis, J., Young, A.C., NISC Comparative Sequencing Program, Bouffard, G.G., Blakesley, R.W., Murray, P.R., et al. (2009). Topographical and temporal diversity of the human skin microbiome. Science 324, 1190–1192.

73. Schetelig, M.F., Lee, K.-Z., Otto, S., Talmann, L., Stökl, J., Degenkolb, T., Vilcinskas, A., and Halitschke, R. (2018). Environmentally sustainable pest control options for Drosophila suzukii. J. Appl. Entomol. 142, 3–17.

74. Cloonan, K.R., Abraham, J., Angeli, S., Syed, Z., and Rodriguez-Saona, C. (2018). Advances in the chemical ecology of the spotted wing drosophila (Drosophila suzukii) and its applications. J. Chem. Ecol. 44, 922–939.

75. Mazzetto, F., Gonella, E., Crotti, E., Vacchini, V., Syrpas, M., Pontini, M., Mangelinckx, S., Daffonchio, D., and Alma, A. (2016). Olfactory attraction of Drosophila suzukii by symbiotic acetic acid bacteria. J. Pest Sci. 89, 783–792.

76. Allgood, D.W., and Yee, D.A. (2017). Oviposition preference and offspring performance in container breeding mosquitoes: evaluating the effects of organic compounds and laboratory colonisation. Ecol. Entomol. 42, 506–516.

77. Diepenbrock, L.M., Swoboda-Bhattarai, K.A., and Burrack, H.J. (2016). Ovipositional preference, fidelity, and fitness of Drosophila suzukii in a co-occurring crop and non-crop host system. J. Pest Sci. 89, 761–769.

78. Jaenike, J. (1978). On optimal oviposition behavior in phytophagous insects. Theor. Popul. Biol. 14, 350–356.

79. Valladares, G., and Lawton, J.H. (1991). Host-plant selection in the holly leaf-miner: Does mother know best? J. Anim. Ecol. 60, 227– 240.

80. Bentley, M.D., and Day, J.F. (1989). Chemical ecology and behavioral aspects of mosquito oviposition. Annu. Rev. Entomol. 34, 401–421.

81. Lampman, R.L., and Novak, R.J. (1996). Oviposition preferences of Culex pipiens and Culex restuans for infusion-baited traps. J. Am. Mosq. Control Assoc. 12, 23–32.

82. Clements, A. (1999). The Biology of Mosquitoes Volume 2: Sensory Reception and Behaviour (CABI Publishing).

83. Wood, D.M., Dang, P.T., and Ellis, R.A. (1979). The Insects and Arachnids of Canada Part 6: The Mosquitoes of Canada - Diptera: Culicidae (Research Branch, Agriculture Canada).

84. Belton, P. (1983). The Mosquitoes of British Columbia (British Columbia Provincial Museum).

85. Carpenter, S., and LaCasse, W. (1955). Mosquitoes of North America (North of Mexico) (University of California Press).

86. Eneh, L.K., Fillinger, U., Borg Karlson, A.K., Kuttuva Rajarao, G., and Lindh, J. (2019). Anopheles arabiensis oviposition site selection in response to habitat persistence and associated physicochemical parameters, bacteria and volatile profiles. Med. Vet. Entomol. 33, 56–67.

87. Schulz-Bohm, K., Martín-Sánchez, L., and Garbeva, P. (2017). Microbial volatiles: Small molecules with an important role in intra- and inter-kingdom interactions. Front. Microbiol. 8, 2484.

88. Ezra, D., and Strobel, G.A. (2003). Effect of substrate on the bioactivity of volatile antimicrobials produced by Muscodor albus. Plant Sci. 165, 1229–1238.

89. Du, Y., and Millar, J.G. (1999). Electroantennogram and oviposition bioassay responses of Culex quinquefasciatus and Culex tarsalis (Diptera: Culicidae) to chemicals in odors from Bermuda grass infusions. J. Med. Entomol. 36, 158–166.

90. Millar, J.G., Chaney, J.D., and Mulla, M.S. (1992). Identification of oviposition attractants for Culex quinquefasciatus from fermented Bermuda grass infusions. J. Am. Mosq. Control Assoc. 8, 11–17.

91. Meijerink, J., Braks, M.A.H., Brack, A.A., Adam, W., Dekker, T., Posthumus, M.A., van Beek, T.A., and van Loon, J.J.A. (2000). Identification of olfactory stimulants for Anopheles gambiae from human sweat samples. J. Chem. Ecol. 26, 1367–1382.

92. Lee, J.-H., Wood, T.K., and Lee, J. (2015). Roles of indole as an interspecies and interkingdom signaling molecule. Trends Microbiol. 23, 707–718.

93. Brodie, B., Gries, R., Martins, A., VanLaerhoven, S., and Gries, G. (2014). Bimodal cue complex signifies suitable oviposition sites to gravid females of the common green bottle fly. Entomol. Exp. Appl. 153, 114–127.

94. Krupke, C.H., Roitberg, B.D., and Judd, G.J.R. (2002). Field and laboratory responses of male codling moth (Lepidoptera: Tortricidae) to a pheromone-based attract-and-kill strategy. Environ. Entomol. 31, 189–197.

95. Suckling, D.M. (2000). Issues affecting the use of pheromones and other semiochemicals in orchards. Crop Prot. 19, 677–683.

96. Haverkamp, A., Hansson, B.S., and Knaden, M. (2018). Combinatorial codes and labeled lines: How insects use olfactory cues to find and judge food, mates, and oviposition sites in complex environments. Front. Physiol. 9, 49.

97. Riffell, J.A., Shlizerman, E., Sanders, E., Abrell, L., Medina, B., Hinterwirth, A.J., and Kutz, J.N. (2014). Flower discrimination by pollinators in a dynamic chemical environment. Science 344, 1515– 1518.

98. Sachs, J., and Malaney, P. (2002). The economic and social burden of malaria. Nature 415, 680–685.

99. Costa-da-Silva, A.L., Navarrete, F.R., Salvador, F.S., Karina-Costa, M., Ioshino, R.S., Azevedo, D.S., Rocha, D.R., Romano, C.M., and Capurro, M.L. (2013). Glytube: A conical tube and Parafilm M-based method as a simplified device to artificially blood-feed the dengue vector mosquito, Aedes aegypti. PLoS ONE 8, e53816.

100. Byrne, K.J., Gore, W.E., Pearce, G.T., and Silverstein, R.M. (1975). Porapak-Q collection of airborne organic compounds serving as models for insect pheromones. J. Chem. Ecol. 1, 1–7.

101. van Den Dool, H., and Kratz, P. (1963). A generalization of the retention index system including linear temperature programmed gas-liquid partition chromatography. J. Chromatogr. A 11, 463–471.

102. Thiel, V., Brinkhoff, T., Dickschat, J.S., Wickel, S., Grunenberg, J., Wagner-Döbler, I., Simon, M., and Schulz, S. (2010). Identification and biosynthesis of tropone derivatives and sulfur volatiles produced by bacteria of the marine Roseobacter clade. Org Biomol Chem 8, 234–246.

